# The assembly of microbial communities on red sandstone surfaces was shaped by dispersal limitation and heterogeneous selection

**DOI:** 10.1101/2024.10.02.616369

**Authors:** Bowen Wang, Chengshuai Zhu, Xin Wang, Tianyu Yang, Bingjian Zhang, Yulan Hu

## Abstract

Understanding the role of microbiota on stone surface is essential for developing effective grottoes conservation strategies. However, ecological feature of microbial communities on stone surface have been rarely investigated systematically. In this study, we explored diversity, assembly, and functional profiles of microbial communities on the red sandstone surface of the Leshan Giant Buddha from microbial ecology perspective. The results show that Proteobacteria, Actinobacteria, Cyanobacteria, and Ascomycota are the dominant phyla. Fundamental metabolic pathways are maintained during the formation of visually distinguishable microbial communities, but gene profiles vary across microbial communities of different colors. Ecological modelling suggests that selective pressure from the harsh stone surface environment fostered the interplay of dispersal limitation and heterogeneous selection during community assembly. The assembly of visually distinct microbial communities is linked to a narrower ecological niche, higher proportion of habitat specialists, and sparser network structure. Microbial-mediated ammonium assimilation and nitrogen mineralization might be the two prominent processes that contribute to stone biodeterioration. This study deepens our understanding of the assembly mechanisms and functional potentials of microbial communities on stone cultural heritage surfaces, provides microbial ecological insights for the conservation of these cultural treasures.

**Importance:** Minimal systematically research on the ecological interpretation of stone biodeterioration. This study reports dispersal limitation and heterogeneous selection shape the microbial community assembly responsible for the biodeterioration of red sandstone. Furthermore, fundamental metabolic processes of microbial communities, such as ammonium assimilation and nitrogen mineralization, are identified as contributors to stone biodeterioration. This study improves our understanding of microbial community assembly and their functional roles, providing a microbial ecological basis for developing effective strategies for the conservation of stone cultural heritage.

## 1. Introduction

Grottoes are a common form of stone cultural heritage. The stone surface of grottoes are integral components of ecosystems, providing critical habitats for microbial communities, serving as crucial habitats for microorganisms organized into diverse microbial communities^1–3^. Under specific conditions, these microbial communities can reinforce the stone through metabolic processes like biomineralization^1,4–7^. However, in most cases, the microbial communities on stone surface not only diminish the aesthetic value of the artifacts by obscuring and deteriorating the carvings but may also, in some cases, compromise the structural integrity of the grottoes through severe erosion processes^3,8–11^. Therefore, cleaning and controlling the formation of microbial communities on stone surface are essential tasks in the conservation of stone cultural heritage. Understanding the characteristics of microbial communities on stone surface is crucial for developing effective strategies to mitigate microbial deterioration. Currently, research in this field primarily focuses on the taxonomic diversity of microorganisms and their biogeochemical cycling and metabolic capabilities. These studies have revealed the key microbial groups involved in the biodeterioration of stone cultural heritage and their associated biogeochemical processes and metabolic responses^10,12–17^. However, ecological understanding of how microbial communities assemble on the stone surface remains significantly limited.

The ecological mechanisms that govern microbial community assembly are one of the main scientific questions in ecological research^18–20^. It is generally believed that the assembly of local microbial communities is controlled by the combined effects of deterministic and stochastic processes^21–23^. Deterministic processes are used to describe the traditional niche-based theory, which hypothesize that species traits, interspecific interactions, and environmental factors govern community structure^24,25^. In contrast, stochastic processes are based on the neutral theory, which illustrate that community assembly is driven by stochastic events such as birth, death, colonization, extinction, and speciation^26,27^. Another quantitative framework for understanding ecological interactions and community assembly is the dissimilarity-overlap curve (DOC), which describes how community dissimilarity changes with the overlap of shared taxa^28^. The niche on the stone surface exposes microorganisms to environmental stresses (abiotic and biotic constraints that limit species productivity and ecosystem development^29^) from nutrient and water scarcity, as well as extreme fluctuations in temperature, humidity, and solar UV radiation^1^. Generally, environmental stress increases deterministic assembly while reducing stochastic assembly, because species with higher tolerance or adaptability to these stressors tend to survive and thrive^29–31^. Furthermore, under high-stress conditions, microbial dispersal limitations are enhanced because elevated stress impedes colonization and successful establishment^29,32,33^. A recent study on microbial communities on limestone surface revealed that bacteria and fungi in this environment are primarily dominated by the mutual effect of dispersal limitation (stochastic processes) and homogeneous selection (deterministic processes)^3^. However, it is currently unknown whether such community assembly mechanisms are specific to certain stone type, geographic location, or influenced by other factors.

The Leshan Giant Buddha, a world-renowned UNESCO World Heritage site, is a typical example of red sandstone grotto cultural heritage. Previous studies found abundant nitrogen-metabolizing microorganisms and ammonia monooxygenase genes, linked to biodeterioration, on the surface of the Leshan Giant Buddha, which suggests that the deterioration of the stone is closely related to microbial growth^34–36^. The cleaning of stone surface is a crucial task for conservation of the grotto. The current issue is that despite regular cleaning of the stone surface by heritage conservators, visually distinguishable microbial communities persistently reappear after each cleaning. However, our understanding of how microorganisms assemble microbial communities on the stone surface of the Leshan Giant Buddha, as well as on other red sandstone surfaces, remains limited. There is an essential need to investigate the mechanisms by which microbes survive and how they form microbial communities within such ecosystems under various environmental stresses from the perspective of microbial ecology. Therefore, this study uses the Leshan Giant Buddha, a UNESCO World Heritage site, as a case study to provide a comprehensively microbial ecological evaluation to the microbial communities on the surface of this red sandstone cultural heritage. The goal of this study is to examine whether the ecological characteristics of microbial communities on red sandstone surfaces are associated with adaptation to harsh environmental conditions. Specifically, we investigate how these microorganisms assemble into stable communities under repeated wet-dry cycles, nutrient limitations, and UV radiation, and what functional capacities they exhibit. Additionally, in this study, we also performed network and niche metrics analysis to better understand the assembly of microbial communities on the red sandstone surface. Network analysis reveals co-occurrence patterns and potential interactions among microbial taxa, while niche metrics such as niche breadth and overlap characterize habitat specialization and resource utilization. These niche-based analyses further allow us to distinguish between generalist (thrive across a wide range of conditions) and specialist (restricted to narrower ecological niches). Our focus extends beyond the interactions within the microbiota to include the interactions among microorganisms, environmental factors, and the stone substrate.

## 2. Results

### 2.1. Taxonomic composition varies by the color of microbial communities

Samples of three different types of visually distinguishable microbial communities, along with blank samples, were collected from the stone surface of the Leshan Giant Buddha. In general, the total 27 samples were grouped and named according to their respective colors (Bla, black; Gre, green; Whi, white; and CK, blank control) (Fig. 1). We then performed amplicon sequencing of conserved phylogenetic marker genes (16S and ITS rRNA) to identify bacteria and fungi within the microbial communities. After quality filtering and rarefaction, a total of 8,791 and 2,404 non-zero ASVs were obtained from the 16S and ITS sequences, respectively. Each group of data overlap with others, but they are not entirely identical or inclusive of each other (Fig. S1).

**Figure 1.**
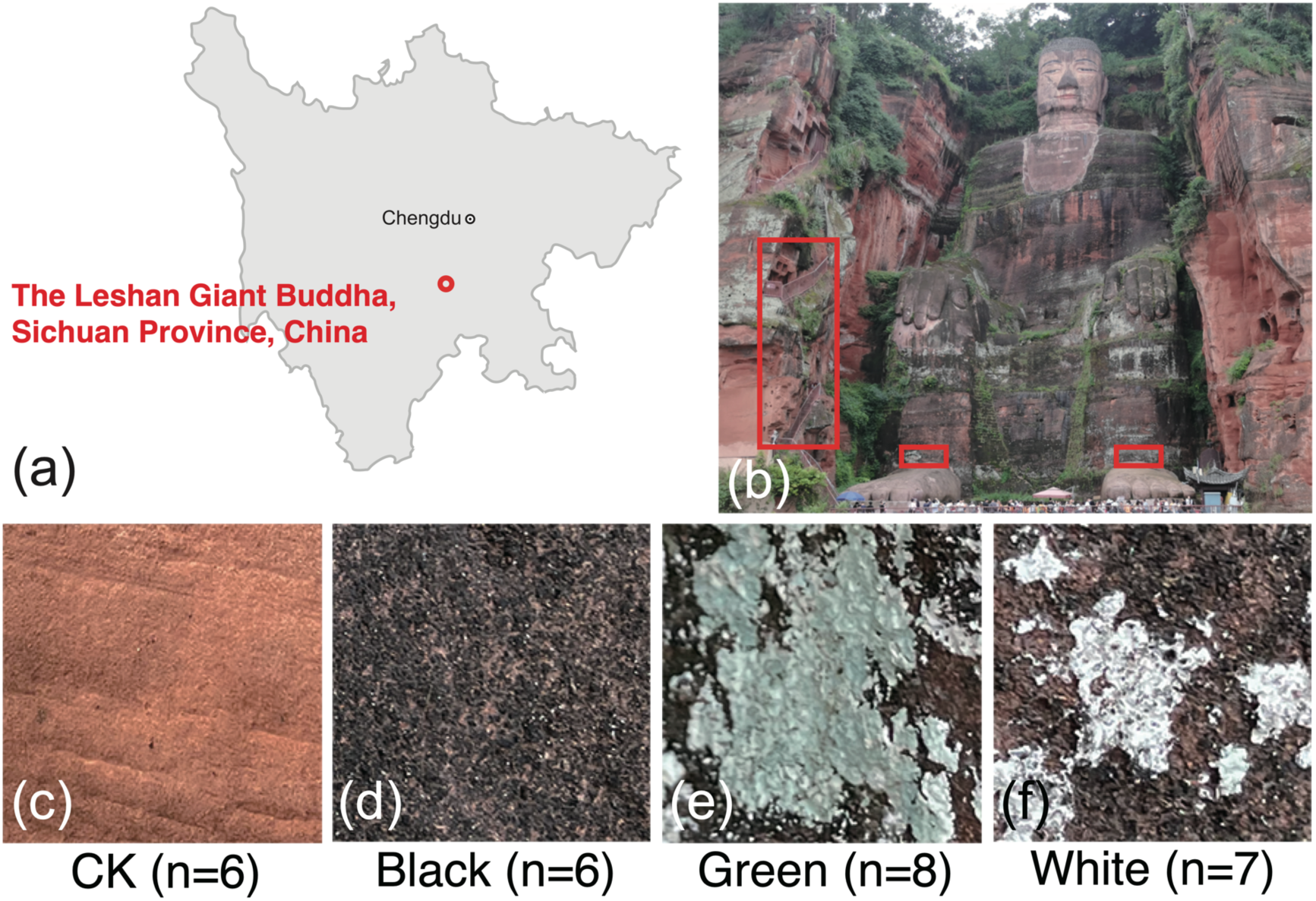
Representative examples and sampling of microbial communities on the red sandstone surface of the Leshan Giant Buddha. (a). Location of the Leshan Giant Buddha. (b). Sampling regions of the samples. Red boxes here indicate the specific sampling regions. (c). Representative examples of CK samples. (d). Representative examples of Black samples. (e). Representative examples of Green samples. (f). Representative examples of White samples. Notes: “n” in picture c-f represents the number of samples used in this study within each group.

To get taxonomic insights into the microbiota, we annotated the sequencing data, then visualized top 10 dominant phyla, top 20 dominant genus of bacteria (Fig. 2 a; Fig. S2 a; Table S1, S2) and fungi (Fig. 2 b; Fig. S2 b; Table S3, S4) across all samples. The results show that among all samples, Proteobacteria, Actinobacteria, and Cyanobacteria are the dominant top three phyla in bacteria, comprising at least 70% of the total bacteria in each sample. Ascomycota is the most dominant phylum in fungi, comprising at least 50% of the total fungi in each sample.

**Figure 2.**
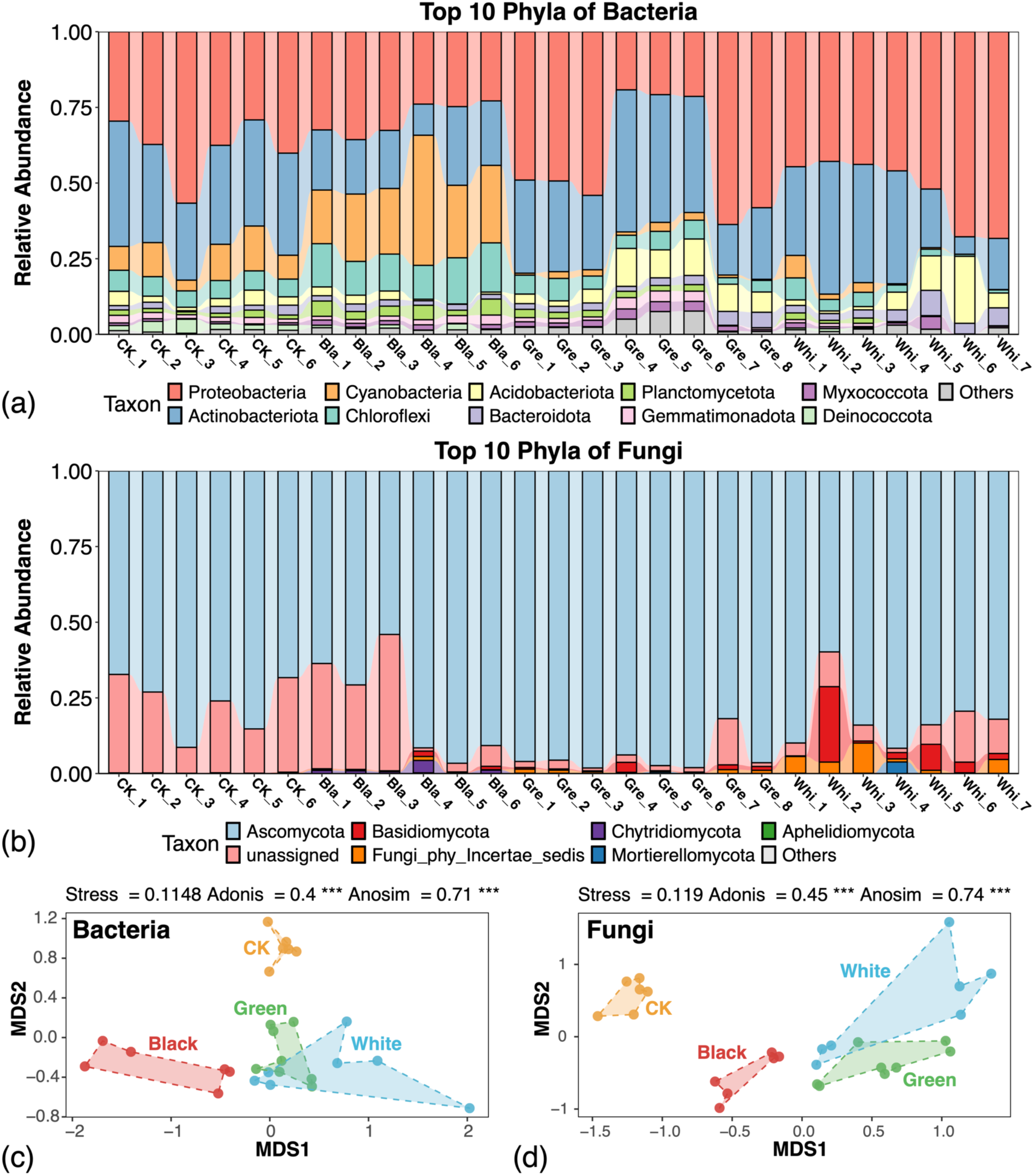
Taxonomic composition and distinct taxonomic structures of the microbial communities. (a). Relative abundances of dominant bacteria among all samples in phylum level. (b). Relative abundances of dominant fungi among all samples in genus level. (c). Non-metric multidimensional scaling (NMDS) ordination based on Bray-Curtis dissimilarities at the ASV level for bacteria. (d). Non-metric multidimensional scaling ordination based on Bray-Curtis dissimilarities at the ASV level for fungi. Notes: The triple asterisks *** represents the P < 0.001.

To estimate the separation pattern between each group of microbial communities, non-metric multidimensional scaling (NMDS) based on Bray-Curtis dissimilarity at the ASV level was performed and then conducted Anosim (analysis of similarities) and PERMANOVA (*i.e.*, adonis) tests (Fig. 2 e, f). The results show that both bacteria and fungi among the four groups of samples reveal significant and major differences (P < 0.001). In summary, the structure of microbial communities varies according to the coloration of the samples, which indicates distinct microbial compositions among samples of different colors.

### 2.2. Dispersal limitation and heterogeneous selection dominants the assembly of red sandstone surface microbial communities

To characterize the assembly mechanism of the red sandstone surface microbial communities, we applied null model analysis^37,38^ and DOC model analysis (Fig. 3)^28^. Null model analysis showed that both deterministic and stochastic processes controlled the assembly of microbial communities, but the relative importance of the two processes in community assembly vary among sample types (Fig. 3 a, b). In general, dispersal limitation dominated the assembly of both bacteria communities (60.11%) and fungi communities (55.56%) on stone surface. Bacteria revealed overall elevated deterministic processes than fungi. Heterogeneous selection and dispersal limitation can explain about 91% of the assembly mechanism of the bacteria communities. While heterogeneous selection and dispersal limitation were also main processes for the assembly of fungi communities; drift played a more pivotal role in the assembly processes of fungi communities (20.23%) than in the bacteria ones (5.41%). Bacteria samples revealed significantly higher value of betaNTI than fungi ones (Fig. 3 c). Significant elevation of betaNTI was detected among White groups of bacteria samples comparing with CK (Fig. 3 d). By contrast, significant elevation of betaNTI was detected among Black as well as Green groups of fungi samples comparing with CK (Fig. 3 e). The DOC model analysis demonstrated a significantly negative correlation between overlap and dissimilarity in both bacteria and fungi communities (P < 0.001), which indicated a general dynamic and interspecific interaction within these microbial communities (Fig. 3 f, g). The fitted slope (Fns) was slightly higher in bacteria (0.3351) than in fungi (0.2691), which suggests that bacterial communities exhibited more rapid compositional turnover. These complex dynamic interactions among microorganisms in the microbiota might lead to competition within the shared niche for survival.

**Figure 3.**
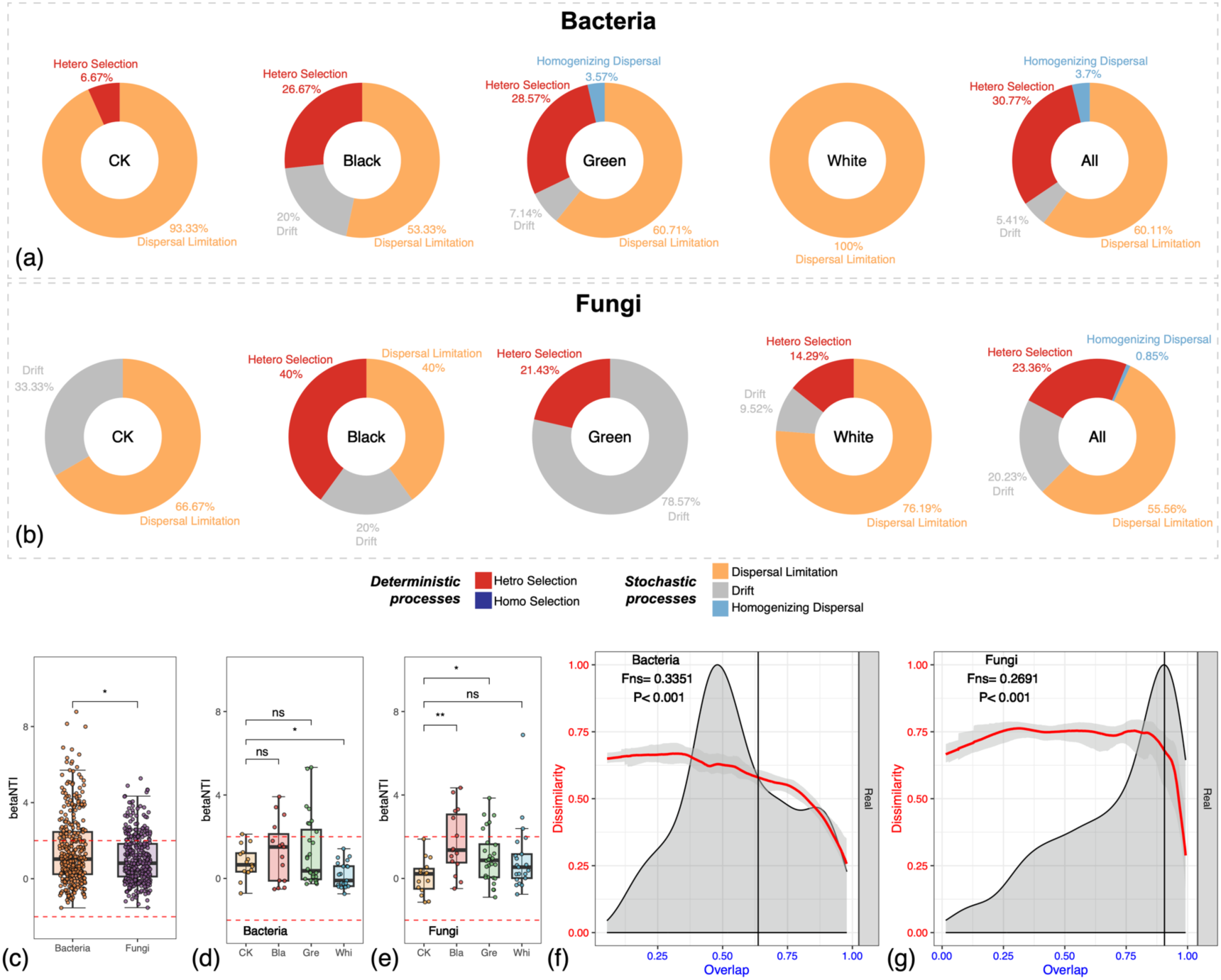
Assembly dynamics of the red sandstone surface microbial communities. (a). The relative importance of assembly processes in bacteria communities. (b). The relative importance of assembly processes in fungi communities. (c). The differences of betaNTI between bacteria and fungi communities. (d). Changes of betaNTI among three groups of bacteria samples comparing with CK. (e). Changes of betaNTI among three groups of fungi samples comparing with CK. (f). Dissimilarity-overlap curve (DOC) analysis of the universality of bacteria dynamics among samples. (g). DOC analysis of the universality of fungi dynamics among samples.

### 2.3. Specialization prevails over flexibility in microbial community formation on red sandstone surface

To delve deeper into the dynamic interactions among microorganisms within the microbiota, we further investigated the niche characterization of microbial communities on red sandstone surface. According to Levins niche breadth theory, the distribution of microbiota composition can be used as an indicator of the environmental conditions^39^. Here, we calculated niche breadth (*Bcom*) and niche overlap (*Ocom*) as indicators to describe the habitat niche characterization of both bacteria and fungi making up the stone surface microbial communities. First, the *Bcom* was calculated with Levins index (Fig. 4 a-c). The results showed that the visually distinguishable microbial communities revealed decreased overall *Bcom* comparing with CK group. Such results were also further validated with Shannon index (Fig. S3 a-c). Then, the *Ocom* was calculated with Schoener index. The *Ocom* for both bacteria and fungi displayed a similar trend that CK samples revealed the highest *Ocom*, followed by Black, Green, and White, in descending order. All pairwise comparisons were statistically significant, indicating distinct niche overlaps among the different samples. When comparing *Ocom* across all samples, bacteria revealed a significantly higher *Ocom* than fungi (Fig. 4 d-f). Such results were also further validated with Pianka index (Fig. S3 d-f). In summary, both bacteria and fungi communities exhibit significant variations in *Bcom* and *Ocom* across different sample types (CK, Black, Green, White). In general, CK samples displayed widest niche breadth and highest overlap level compared with the rest groups of samples. These results suggested that the formation of visually distinguishable microbial communities on stone surface requires species to be more specialized, rather than relying on diverse and flexible species with metabolic flexibility and redundancy.

**Figure 4.**
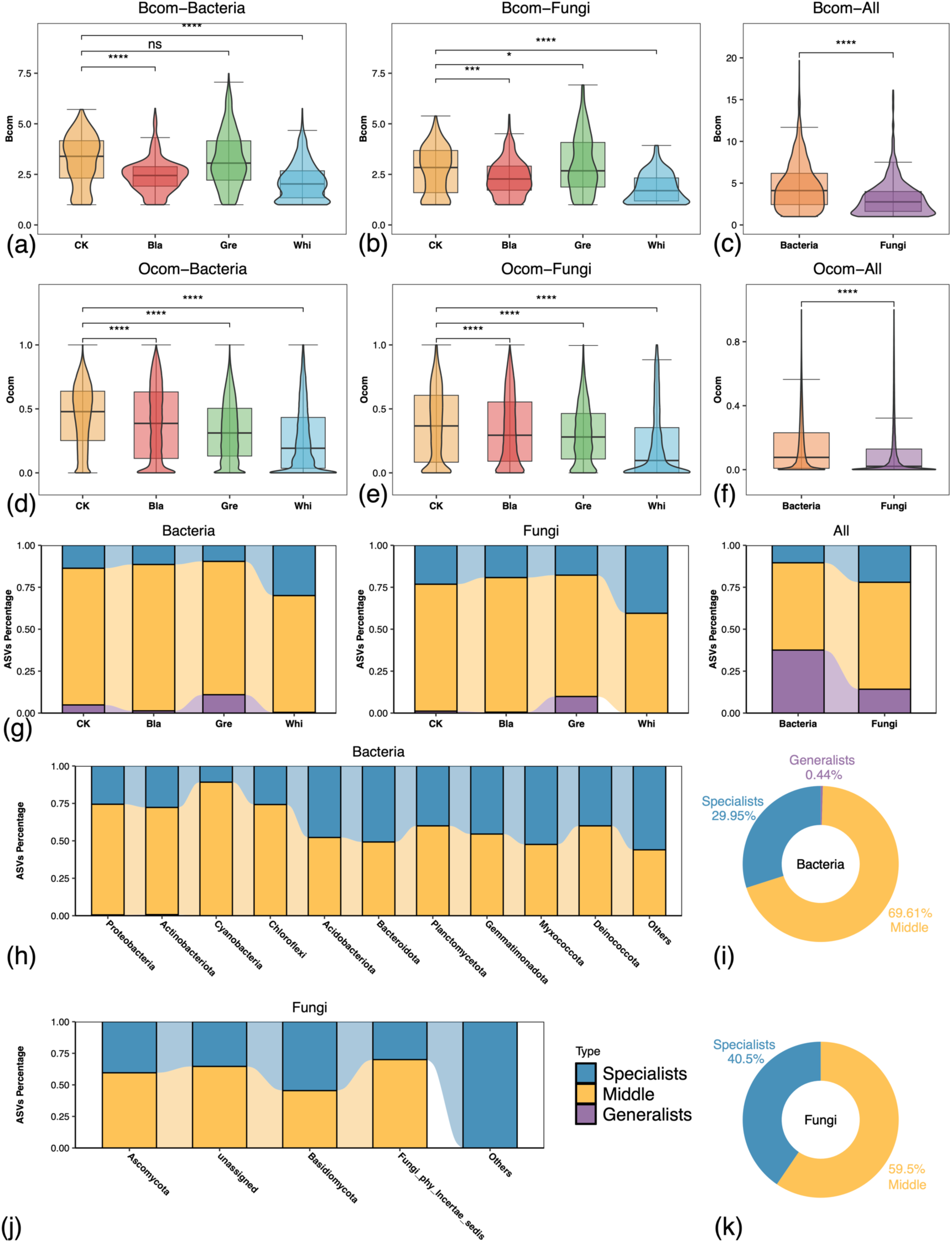
Niche characterization of the red sandstone surface microbial communities based on Levins niche breadth index and Schoener niche overlap index. (a). Average habitat niche breadth (*Bcom*) differences among three groups of bacteria samples comparing with CK in bacteria. (b). *Bcom* differences among three groups of fungi samples comparing with CK in fungi. (c). *Bcom* differences between bacteria and fungi. (d). Average habitat niche breadth (*Ocom*) differences among three groups of bacteria samples comparing with CK in bacteria. (e). Average *Ocom* differences among three groups of fungi samples comparing with CK in fungi. (f). Average *Ocom* differences between bacteria and fungi. (g). Proportional distribution of specialists, middle, and generalists in each sample group, and between bacteria and fungi. (h). Proportional distribution of specialists, middle, and generalists of bacteria at phylum level based on Levins niche breadth index. (i). Overall proportion of the habitat niche breadth of bacteria at phylum level. (j). Proportional distribution of specialists, middle, and generalists of fungi at phylum level. (k). Overall proportion of the habitat niche breadth of fungi at phylum level. Notes: The *Bocm* was calculated with Levins index. The *Ocom* was calculated with Schoener index. All ASVs were defined as specialists, middle or generalists according to Levins niche breadth index. The asterisks in the boxplots stand for the significance of the Wilcoxon signed-rank test. The “ns” stands for no significance, single asterisk stands for P < 0.05, double asterisks stand for P < 0.01, Triple asterisks stand for P < 0.001, Four asterisks stand for P < 0.0001.

To further explore the inference, we also calculated the proportion of generalists and specialists among the microbial communities (Fig. 4 g-k). Generalists are species that are better suited to survive in diverse and challenging environments and often play pivotal roles within their communities. Specialists, on the other hand, have a more limited range of habitats and are specifically adapted to environmental conditions^40,41^. The results showed that at the ASVs level, the proportion of specialists in each type of sample was higher than that of generalists in both bacteria and fungi (Fig. 4 g). To determine whether there were differences between all bacteria and fungi, we also compared the proportion of specialists and generalists within these groups. The results showed that in each group of both bacteria and fungi, the proportion of specialists was higher than that of generalists. But, bacteria revealed overall higher proportion of generalists than that of specialists (Fig. 4 g). At the phylum level, up to 69.61 % of the bacteria were classified as the middle; the proportion of specialists (29.95 %) was higher than that of generalists (0.44 %) (Fig. 4 i, Table S5). Myxococcota revealed the largest fraction of specialists in bacteria (52.4%) among the top 10 dominant phyla. (Fig. 4 h, Table S5). For the fungi, 59.5% were classified as middle, 40.5% as specialists, and none of the ASV is classified as generalists. (Fig. 4 k, Table S6). This suggested that specialists prevail over generalists in the dominant phyla within the assembly of microbial communities. Notably, the non-dominant phyla (others) of both bacteria and fungi exhibited the highest proportion of specialists (Fig. 4 h, j; Table S5, S6). This suggested that the non-dominant phyla within the community might have even lower metabolic flexibility compared to the dominant phyla.

### 2.4. Sparser network structure during the formation of visually distinguishable microbial communities on red sandstone surface

Given the non-random assembly patterns observed in red sandstone surface microbial communities, we proceeded to construct microbial co-occurrence networks of both bacteria and fungi in different types of microbial communities to further explore the interaction patterns within these communities (Fig. 5). Subnetworks corresponding to each group were extracted from the global network constructed using all samples to obtain more robust group-specific networks.

**Figure 5.**
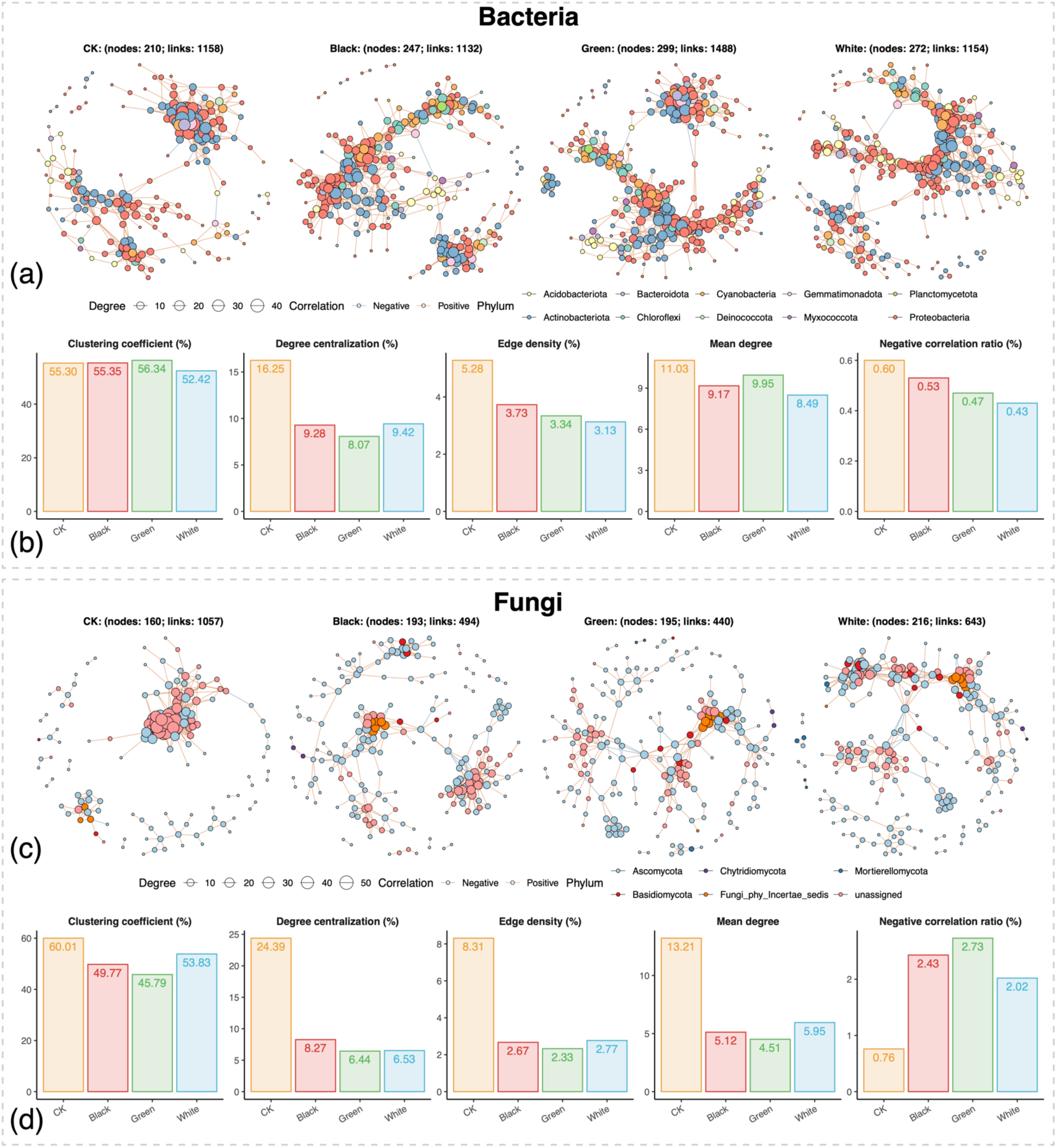
Topological features of the co-occurrence network within microbial communities on the stone surface. (a). Co-occurrence network of bacterial communities. (b). Topological comparisons of the four bacterial subnetworks. (c). Co-occurrence network of fungal communities. (d). Topological comparisons of the four fungal subnetworks. Notes: Subnetworks corresponding to each group were extracted from the global network constructed using all samples to obtain more robust group-specific networks.

Topological feature analysis revealed significant differences in bacterial networks among the different community types (Fig. 5a). Compared with the CK group, the three visually distinguishable microbial communities exhibited higher numbers of nodes and edges, which indicated that their formation was accompanied by a more complex bacterial association structure (Fig. 5a). However, in terms of structural parameters, the CK group showed higher clustering coefficient, degree centralization, edge density, and mean degree than the other three groups (Fig. 5b), which suggests that the visually distinguishable communities possessed a sparser local connectivity and overall network structure. Meanwhile, all three visually distinguishable communities exhibited lower proportions of negative correlations than CK, which implies that competitive or exclusionary interactions among bacterial members were reduced and replaced by more cooperative associations. The fungal networks showed a similar trend to the bacterial ones. The fungal networks of the three visually distinguishable communities underwent marked structural simplification, with fewer edges and reduced network density, which suggested weakened associations among fungal taxa (Fig. 5c, d). At the same time, the Black and Green groups displayed higher proportions of negative correlations than CK, which indicated enhanced antagonistic interactions among fungal members within the visually distinguishable communities (Fig. 5d). Overall, compared with CK, the microbial networks of the visually distinguishable communities became notably sparser in their connectivity.

### 2.5. Metabolic potential and nitrogen cycling of microbial communities on red sandstone surface

To elucidate the metabolic processes that enable the survival of microbial communities on the stone surface, we performed metagenomic sequencing for each type of microbial communities. Shotgun metagenomic sequencing generated approximately 9.6-13.2 Gb of raw data per sample. After quality control, 6.4-8.8 Gb of high-quality clean reads, comprising 206,491,166 clean reads in total, were retained for downstream analyses (Table S7).

Ten Gene Ontology (GO) terms that could potentially contribute to the assembly of microbial communities on red sandstone surface were identified in all four types of microbial communities^3^. These terms were associated with functions related to organism interactions, microbial survival, and trophic processes. Each of the ten GO terms was detectable in every group (Fig. 6 a).

**Figure 6.**
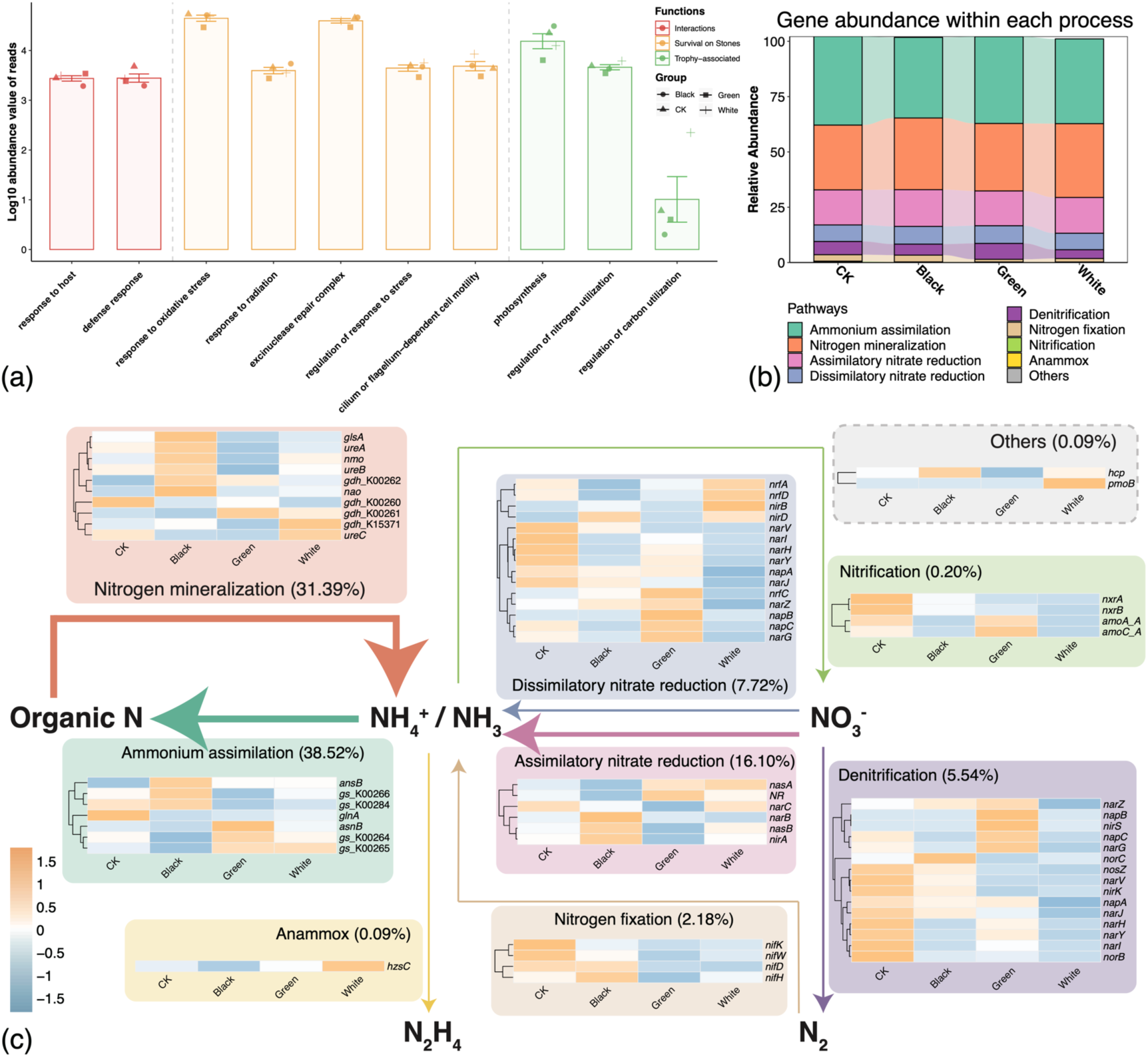
Functional profiles of the stone surface microbiota. (a). GO functional annotation of the stone surface microbiota. (b). Gene abundance within each nitrogen cycle pathway of the stone surface microbiota. (c). Nitrogen cycling potential of the stone surface microbiota. Arrow thickness corresponds to the proportion of functional genes for corresponding metabolic processes.

Nitrogen cycle processes have been broadly reported as closely related with the deterioration of stone^3,9,42–44^. Hence, we performed detailed insight into nitrogen cycle processes. The results showed that nitrogen cycle related genes were widely present across all stone surface microbial communities (Fig. 6 b), indicating that these communities maintain the potential for nitrogen cycling even under extremely nutrient-limited and highly variable environmental conditions. Among all nitrogen cycle pathways, ammonium assimilation and nitrogen mineralization were the dominant processes, which account for about 38.52% and 31.39% of the total abundance of nitrogen cycle related reads, respectively (Fig. 6 c). This finding suggests active nitrogen exchange between microbial communities and the stone substrate. In addition, the relative abundance of nitrogen related genes varied among microbial communities of different color types (Fig. 6 c), which might reflect the effects of local microenvironmental heterogeneity on nutrient availability.

## 3. Discussion

Uncovering the diversity, assembly mechanisms, and functional traits of microbial communities is a central focus in microbial ecology^18,45,46^. Recently, the role of microbial communities on stone surfaces has garnered increasing attention, particularly in the field of stone cultural heritage conservation^47^. In this study, we present a pioneering case study on the ecological roles of microbiota on red sandstone surface, using the Leshan Giant Buddha as an example. The results indicate that the taxonomic composition of microbial communities varies according to their coloration. During the formation of visually distinguishable microbial communities, core metabolic pathways of microbial communities were maintained but gene profiles varied across different types of microbial communities, particularly in nitrogen cycle-related biogeochemical processes. Dispersal limitation and heterogeneous selection dominate the assembly of stone surface microbial communities. Habitat specialists play a more significant role than generalists during the formation of microbial communities on the red sandstone surface. As visible microbial communities form on stone surface, network complexity increases, and resistance to disturbances is enhanced. By comparing our findings with previous studies on microbial communities on stone surface, we inferred the following commonalities and unique characteristics in the ecological features of stone surface microbial communities.

### 3.1. Conservation of core ecological functions under harsh stone surface conditions

In this study, the microbial diversity analysis revealed that Proteobacteria, Actinobacteria, Cyanobacteria, Chloroflexi, and Acidobacteriota dominate the top phyla among the bacteria in all samples, while Ascomycota stands as the most dominant phylum in fungi, despite the distinct microbial community morphology across different sample groups. These phylum-level diversity patterns are largely consistent with previous reports on taxonomic diversity of microbial communities on the surface of other stone types. However, significant differences were observed at the genus level^2–4,35,48–50^. These variations might be attributed to site-specific factors, including the local climate and the type of stone cultural heritage^51^. Comparison with previous studies suggests that core ecological functions of stone surface microbes are conserved across the surfaces of various types of stone. The capability to withstand pressures such as fluctuations between wet and dry conditions, endure UV stress on stone surface, and facilitate ion exchange with the stone appears essential for the persistence of microbial communities on the outdoor stone surface^3,9^.

The nitrogen cycle is likely a conserved and essential pathway that mediates ion exchange between microbial communities and the stone substrate across various types of stone surface ecosystems. This study revealed that distinct microbial communities exhibited similar ecological functional patterns, which highlights the important role of functional redundancy in maintaining the stability of stone surface ecosystems. Dispersal limitation and environmental filtering jointly shaped the taxonomic divergence among communities, while the core nitrogen cycling functions were likely sustained through complementary metabolic strategies. These findings further support that the nitrogen cycle serves as a crucial pathway for ion exchange between microbial communities and the stone substrate, regardless of taxonomic convergence or divergence. For example, previous study reported that the two dominated types of nitrogen cycling genes in limestone surface microbial communities are nitrogen assimilation and mineralization, which is coherent with what we found in red sandstone surface microbial communities in this study^3^. Nitrifying bacteria genera *Nitrospira* and *Nitrosomonas* were significantly enriched in limestone surface microbial communities^3^. In another case study involving regular sandstone, *Nitrosopumilus* and *Nitrospira* were also identified from deteriorated sediments^9^. Furthermore, *Methylobacterium*, another dominant genus identified in our study, comprises species recognized as plant growth-promoting rhizobacteria, with genomes encoding enzymes involved in nitrogen metabolism and ammonia assimilation^52^. Despite the overall functional convergence of nitrogen metabolism, differences in the relative abundance and composition of nitrogen cycling genes were observed among microbial communities of different color types. These variations likely reflect the microenvironmental heterogeneity of the stone surface, where factors such as moisture, organic matter, and light gradients influence nutrient availability and redox conditions^53^. Consequently, environmental filtering and niche specialization may have selected for distinct microbial taxa with complementary nitrogen transformation capabilities. Such taxon-specific metabolic adaptations could explain the observed diversity of nitrogen-related genes while maintaining overall functional redundancy within the system.

Conserved core functions among microbial communities within similar niches are not unique to microbial communities on stone surface. For example, several gut microbiota studies have reported functional, but not compositional, convergence of gut microbiomes^54–57^. Some soil microbiota studies also reported the stable functional structure despite high taxonomic variability^58–60^. In addition, microbial functional genes of ocean microbiota that execute certain functions, such as B_12_ biosynthesis, are widely distributed, while the microbial taxa that carry these genes vary significantly^61^. These results corroborate the ecological hypothesis that environment truly selects for function, rather than taxonomy^62^. Different taxonomic groups could reveal convergent functions as a result of functional redundancy among microbial systems^63^. On stone surface, conserved core functions among various types of stones might be attributed to the selective pressure imposed by the harsh living conditions on the stone surface, favoring the survival of microbial communities with specific functions^58,64–66^.

### 3.2. Assembly of stone surface microbial communities requires a general pattern of dynamic interactions

In this study, we witnessed the extensive microbe-microbe and microbe-environment interactions within the microbial communities on the stone surface. The mutual effect of heterogeneous selection and dispersal limitation controls the assembly processes of both bacterial and fungal communities on the red sandstone surface. Such results differ from previous studies on the assembly processes of microbial communities on limestone surface, where the assembly processes are driven by homogeneous selection and dispersal limitation^3^. This suggests that the assembly mechanisms of microbial communities on stone surface might vary based on stone types, geographical locations, climatic characteristics, or other environmental factors. In addition, the results of the DOC model analysis suggest a general pattern of dynamic interspecific interactions within the communities on the red sandstone surface and high selective pressure on the red sandstone surface^28,67–69^. Such complexity was further testified in this study by the topological feature analysis of the microbial networks. The active biogeochemical metabolism such as ammonium assimilation and nitrogen mineralization processes by the microbial communities also suggests their active interactions with the red sandstone substrate^70^. The elevated complexity of microbial communities might arise from widely functional complementarity and the resulting extensive interactions among microbes^71^.

Here, we propose a hypothesis that a possible model capable of precisely interpreting the assembly processes of stone surface microbial communities is the classic MacArthur consumer-resource model, where species-species interactions are mediated through resource dynamics^72–74^. Because of the nutrient-limited nature of the stone surface, the colonization by pioneer species is primarily driven by deterministic processes linked to nutrient availability, which aligns with commonly observed among natural communities from various locations^75^. Some pioneer species utilize inorganic energy from the stone to establish themselves. These species then leak metabolic by-products into the environment, increasing the diversity of resources and enhancing the biocompatibility of the stone surface. Other species can subsequently utilize these leaked metabolites, in turn releasing new by-products into the environment, further enhancing the biocompatibility of the stone surface, and facilitating the formation of microbial communities. The observed decrease in ecological niche breadth among the visually distinguishable microbial communities in this study also suggests that the utilization of specific resources is more efficient within these communities. We also observed that specialization prevails over flexibility during the formation of microbial communities on red sandstone surface. This is consistent with a previous study on the assembly of microbial communities in a wide range of well-defined resource environments, where habitat specialists also drive the increase of diversity in such microbial communities^19^. Meanwhile, the sparser structure during the formation of the visually distinguishable microbial networks suggests that although the communities are taxonomically diverse, their members are relatively independent or show clear functional differentiation. The reduction in network interactions, especially for fungal communities, indicates that each species occupies a more defined ecological niche within the biofilm, with stronger resource competition or spatial segregation. It can therefore be inferred that different modules within the microbial community might form a micro-ecological isolation, which allows various microbial populations to coexist through spatial and functional differentiation. Such spatial partitioning might occur between the surface and deeper layers of the microbial community. Future research should make more detailed spatial distinctions between surface and subsurface layers and within them to better resolve the mechanisms of community formation. In addition, higher-resolution temporal studies are needed to clarify the dynamic processes underlying microbial community assembly.

### 3.3. Summary

In summary, this study investigated the ecological and functional characteristics of microbial communities on the red sandstone surface of the Leshan Giant Buddha. Under the interplay of dispersal limitation and heterogeneous selection, the microbiota on the red sandstone were assembled into distinct types of visually distinguishable microbial communities, which significantly differed in the taxonomic composition of these communities. However, under the harsh habitat on red sandstone surface, such as UV radiation, fluctuations between dry and wet conditions, and limited nutrient availability, the core functions of the microbe across different types of microbial communities are conserved. These functions are particularly enriched in biogeochemical processes related to nitrogen cycling and responses to environmental stress. The formation of visually distinguishable microbial communities is associated with a narrower niche width, a higher proportion of habitat specialists, and a sparser network structure (Fig. 7). Given the nutrient limitations of the red sandstone surface, the classic MacArthur consumer-resource model might provide a potential framework to further explain the assembly mechanisms of these microbial communities. In the future, in-depth temporal and spatial studies of the ecological characteristics and functional potential of these microbial communities will provide valuable insights into their assembly on red sandstone surfaces. Moreover, microbial deterioration affects not only stone cultural heritage but also a wide range of other cultural heritage materials^76–80^. The microbial ecology-based approach used in this study will help advance research on microbial deterioration mechanisms in other cultural heritage materials, providing a model for future work.

**Figure 7.**
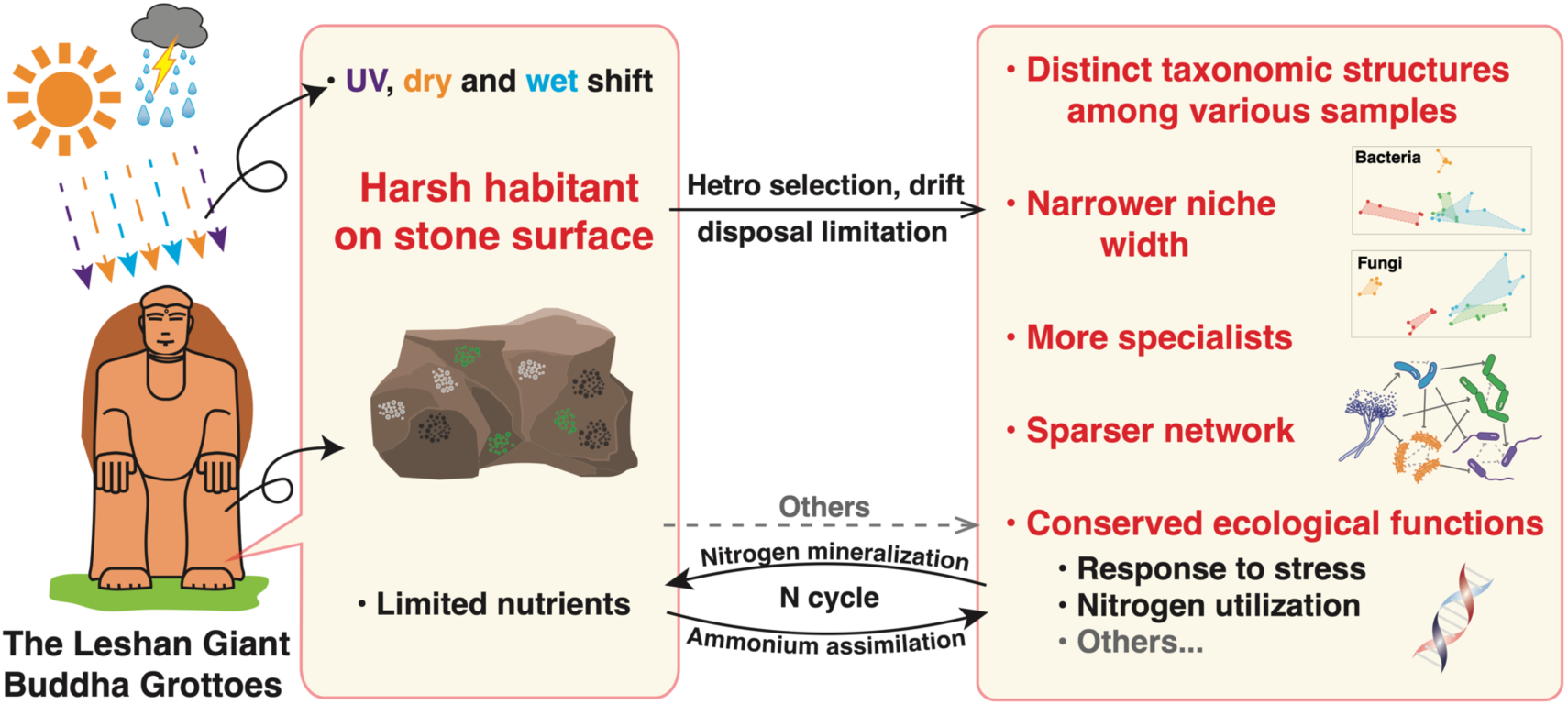
Schematic diagram illustrates the characteristics of microbial communities on the red sandstone surface of the Leshan Giant Buddha

## 4. Materials and Methods

### 4.1. Sampling processes

The Leshan Giant Buddha is a world-famous, typical red sandstone-made, UNESCO World Heritage site. It was built between 713 and 803 during the Tang dynasty, located in the Leshan city, Sichuan province, China. Samples used in this study were collected from the stone surface of the Leshan Giant Buddha between 2019 and 2022. Among these, 6 samples have been used in our previous study^35^. The other 21 samples were newly collected in 2022. Surface samples were scraped using a pre-cleaned steel scalpel and then stored in low temperature until DNA extraction.

### 4.2. DNA extraction and NGS sequencing

The DNA extraction was applied using a genomic DNA extraction kit from TIANGEN (DP302-02) with 0.5 g of sample processed for extraction. The amplicon sequencing was performed as followed. The 16S V3-V4 region rDNA and ITS2 region were amplified by PCR (initial denaturation at 98 ℃ for 30 seconds; 32cycles of denaturation at 98 ℃ for 10 seconds, annealing at 54℃ for 30 seconds, and extension at 72 ℃ for 45 seconds; and then final extension at 72 ℃ for 10 minutes.) with following primers: 0341-F (CCT ACG GGN GGC WGC AG) and 806-R (GAC TAC HVG GGT ATC TAA TCC) for 16S^81^, ITS1FI2 (GTG ART CAT CGA ATC TTT G) and ITS2 (TCC TCC GCT TAT TGA TAT GC) for ITS2. The PCR products were purified by AMPure XT beads (Beckman Coulter Genomics, Danvers, MA, USA) and quantified by Qubit (Invitrogen, USA). The amplicon pools were prepared for sequencing and the size and quantity of the amplicon library were assessed on Agilent 2100 Bioanalyzer (Agilent, USA) and with the Library Quantification Kit for Illumina (Kapa Biosciences, Woburn, MA, USA), respectively, and then sequenced on Illumina NovaSeq 6000 PE250 platform in LC-Bio Technology Co., Ltd, Hang Zhou, Zhejiang Province, China. The raw sequence data were then deposited in the Genome Sequence Archive^82^ in National Genomics Data Center^83^, China National Center for Bioinformation / Beijing Institute of Genomics, Chinese Academy of Sciences (GSA: CRA013382) that are publicly accessible at https://ngdc.cncb.ac.cn/gsa.

The DNA libraries were sequenced on the illumina NovaseqTM 6000 platform by LC Bio Technology CO., Ltd (Hangzhou, China). We are grateful to LC Bio Technology CO., Ltd for assisting in sequencing. DNA concentration was measured using Qubit. Based on the measured concentration, equal-mass DNA samples were pooled from all samples within each group by mixing 100 ng of DNA from each sample. The metagenome sequencing was performed as followed: DNA library was constructed by TruSeq Nano DNA LT Library Preparation Kit (FC-121-4001). DNA was fragmented by dsDNA Fragmentase (NEB, M0348S) by incubate at 37°C for 30min. Library construction begins with fragmented cDNA. Blunt-end DNA fragments are generated using a combination of fill-in reactions and exonuclease activity, and size selection is performed with provided sample purification beads. An A-base is then added to the blunt ends of each strand, preparing them for ligation to the indexed adapters. Each adapter contains a T-base overhang for ligating the adapter to the A-tailed fragmented DNA. These adapters contain the full complement of sequencing primer hybridization sites for single, paired-end, and indexed reads. Single- or dual-index adapters are ligated to the fragments and the ligated products are amplified with PCR. The raw sequence data were then deposited in the Genome Sequence Archive^82^ in National Genomics Data Center^83^, China National Center for Bioinformation / Beijing Institute of Genomics, Chinese Academy of Sciences (GSA: CRA013393) that are publicly accessible at https://ngdc.cncb.ac.cn/gsa.

### 4.3. Taxonomic and functional annotations

For the amplicon sequencing data, the DADA2 R package^84^ in QIIME 2^85^ was used to conduct a complete pipeline to generate raw paired-end fastq sequencing files into merged, denoised, chimera-free sequences and output ASV. The ASV tables for both 16S and ITS were rarified to a minimum sequencing depth (13,622 for 16S and 16,232 for ITS) to mitigate the impact of sequencing depth on subsequent analysis. Representative ASV sequences were classified into different organisms using a naive Bayesian model with RDP classifier^86^, which is based on the SILVA database^87^ (version 138) for 16S or UNITE database^88^ (version 9.0) for ITS.

Metagenome raw sequencing reads were processed to obtain valid reads for further analysis. First, sequencing adapters were removed from sequencing reads using cutadapt (version 1.9). Secondly, low quality reads were trimmed by fqtrim (version 0.94) using a sliding-window algorithm. Window size is 6 bp, and base quality must not be lower than 20. Once quality-filtered reads were obtained, they were de novo assembled to construct the metagenome for each sample by MEGAHIT with the following parameters: *--k-list 21,29,39,59,79,99,119,141 -m 0.8 -t 8 --min-contig-len 200 --continue*. All coding regions (CDS) of metagenomic contigs were predicted by MetaGeneMark (version 3.26). CDS sequences of all samples were clustered by CD-HIT (version 4.6.1)^89^ to obtain unigenes with the following parameters: *-c 0.95 -aS 0.9 -d 0 -g 1 -M 20000 -T 20*. The CDS sequences from all samples are clustered together to group highly similar or identical sequences, and a single representative sequence from each cluster is chosen as a unigene. Unigene abundance for a certain sample were estimated by TPM based on the number of aligned reads by bowtie2 (version 2.2.0)^90^. The TPM value for each Unigene was calculated by dividing its RPK (Reads Per Kilobase) by the sum of RPK values across all Unigenes in the sample, then multiplying the result by 1,000,000 to normalize expression levels to a per-million scale. The functional annotations (GO, KEGG) of unigenes were obtained by aligning them against the NCBI NR database (version 20191121) by DIAMOND (version 0.9.14)^91^ with the following parameters: *-k 25 -e 0.00001 --query-cover 0 --min-score 0 --id 0.* Nitrogen cycling genes were annotated with the NCycDB database^92^. Heatmaps were visualized with pheatmap package in R.

The figures in this study was widely plotted in R project ggplot2 package^93^. Top 10 phyla were visualized in R with ggplot2 and ggalluvial as our previous publications^69,76^. Bray-Curtis distance matrix was calculated in R with Vegan package^94^ (version 2.5.7). NMDS was then generated with Vegan and plotted with ggplot2 package. Adonis and anosim test were used to calculate the significance of β diversity differences.

### 4.4. Microbial communities assembly analyses

The DOC model was used to determine whether microbiota on the red sandstone surface of the Leshan Giant Buddha have unique ecological dynamics in R with DOC package (https://github.com/Russel88/DOC) according to previous reports^3,68,69^. The DOC was obtained by plotting the dissimilarity (Jensen–Shannon divergence) against the overlap fraction of taxa for each pairwise community^28^ with 1000 bootstrap replicates to evaluate the significance and initiation of negative slopes.

Null model analysis was performed as described by Stegen *et al*. ^37,38^ and visualized as previous research^69^. The deviation of phylogenetic and taxonomic diversity was estimated with 999 randomizations. Beta mean nearest taxon distance (βMNTD) and beta nearest taxon index (βNTI) were calculated with picante package in R^95^. Phylogenetic analysis were performed by iqtree2 with 1000 bootstrap^96^. A significant deviation indicated the dominance of deterministic processes, including homogeneous or heterogeneous selection processes when βNTI < −2 or βNTI > 2, respectively. The Bray-Curtis based Raup-Crick (RCbray), the deviation between Bray-Curtis and the null distribution, was then calculated to partition pairwise comparisons that assigned the stochastic processes when |βNTI| < 2. The relative influence of homogenizing dispersal and dispersal limitation was quantified as the fraction of pairwise comparisons with RCbray < −0.95 and RCbray > 0.95, respectively. In addition, dispersal was seen as stochastic processes in this research, not restricted to deterministic or stochastic processes as previously described^3,18^. Low-occurrence ASVs with a prevalence less than 50% and low abundance ASVs with total read counts lower than the number of samples were excluded to ensure the accuracy in taxon-specific assembly characterization and reduce type I errors as presented in previous research^3,69^.

### 4.5. Niche characteristics analysis

Levins niche breadth index, Shannon niche breadth index, Schoener niche overlap index, and Pianka niche overlap index were calculated with spaa package in R^97^. Generalists were defined by ASVs with *Bcom* > 5, whereas specialists were defined by those with *Bcom* < 1.5^3,98^. Significance of the differences was tested by Wilcoxon signed-rank test.

### 4.6. Network construction and topological feature analysis

To obtain more robust group-specific networks, subnetworks corresponding to each group were extracted from the global network constructed using all samples. Microbial co-occurrence networks were constructed according to previous publication^99^ with modifications by ggClusterNet R package^100^. To reduce noise from rare taxa, only the top 400 most abundant ASVs across all samples were retained for network analysis. Pairwise correlations between ASVs were calculated using the Spearman’s rank correlation coefficient. A correlation coefficient threshold of |r| > 0.75 and P values < 0.001 were retained for network construction to ensure significant and robust associations. Group-specific subnetworks were derived by retaining ASVs with nonzero abundance that appeared in the global network and were not isolated within the extracted subnetwork, along with their edges. The subnetworks were then visualized with ggraph in R^101^. Network topology metrics of each subnetwork were computed with the igraph package in R^102^.

## 4.7 Acknowledgements of English edition

During the preparation of this work the authors used ChatGPT in order to check the English grammar. After using this tool, the authors reviewed and edited the content as needed and take full responsibility for the content of the publication.

## Availability of data and materials

The raw sequence data reported in this paper have been deposited in the Genome Sequence Archive in National Genomics Data Center, China National Center for Bioinformation / Beijing Institute of Genomics, Chinese Academy of Sciences (GSA: CRA013382 & GSA: CRA013393) that are publicly accessible at https://ngdc.cncb.ac.cn/gsa.

All scripts used in this study are publicly available at the GitHub repository (https://github.com/Bowenw6/Microbiome_Buddha_Grotto_2023).

## Author Contributions Statement

**Bowen Wang**: Conceptualization, Data curation, Formal analysis, Investigation, Methodology, Resources, Visualization, Writing - Original Draft, Writing - Review & Editing. **Chengshuai Zhu**: Conceptualization, Data curation, Investigation, Resources. **Xin Wang**: Conceptualization, Investigation, Resources. **Tianyu Yang**: Conceptualization, Investigation, Resources. **Bingjian Zhang**: Conceptualization, Funding acquisition, Supervision. **Yulan Hu**: Conceptualization, Funding acquisition, Project administration, Supervision, Writing - Review & Editing.

## References

1 Liu, X. B., Koestler, R. J., Warscheid, T., Katayama, Y. & Gu, J. D. Microbial deterioration and sustainable conservation of stone monuments and buildings. Nat Sustain 3, 991–1004 (2020). 10.1038/s41893-020-00602-5

2 Brewer, T. E. & Fierer, N. Tales from the tomb: the microbial ecology of exposed rock surfaces. Environ Microbiol 20, 958–970 (2018). 10.1111/1462-2920.14024

3 He, J. et al. From surviving to thriving, the assembly processes of microbial communities in stone biodeterioration: A case study of the West Lake UNESCO World Heritage area in China. Sci Total Environ 805, 150395 (2022). 10.1016/j.scitotenv.2021.150395

4 Qiang, L., Zhang, B. B., Yang, B. X. & Ged, C. Q. Deterioration-Associated Microbiome of Stone Monuments: Structure, Variation, and Assembly. Appl Environ Microbiol 84, e02680-02617-02617 (2018).

5 Li, T., Hu, Y. & Zhang, B. Biomineralization Induced by Colletotrichum acutatum: A Potential Strategy for Cultural Relic Bioprotection. Front Microbiol 9, 1884 (2018).

6 Jroundi, F., Gonzalez-Muñoz, M. T. & Rodriguez-Navarro, C. in Microorganisms in the Deterioration and Preservation of Cultural Heritage (ed Edith Joseph) 281–299 (Springer International Publishing, 2021).

7 Liu, X. et al. Biofilms on stone monuments: biodeterioration or bioprotection? Trends Microbiol 30, 816–819 (2022). 10.1016/j.tim.2022.05.012

8 Li, J. et al. The active microbes and biochemical processes contributing to deterioration of Angkor sandstone monuments under the tropical climate in Cambodia – A review. J Cult Herit 47, 218–226 (2021). 10.1016/j.culher.2020.10.010

9 Ding, X. et al. Microbiome and nitrate removal processes by microorganisms on the ancient Preah Vihear temple of Cambodia revealed by metagenomics and N-15 isotope analyses. Appl Microbiol Biotechnol 104, 9823–9837 (2020). 10.1007/s00253-020-10886-4

10 Wu, F. et al. Metagenomic and metaproteomic insights into the microbiome and the key geobiochemical potentials on the sandstone of rock-hewn Beishiku Temple in Northwest China. Sci Total Environ 893, 164616 (2023). 10.1016/j.scitotenv.2023.164616

11 Fidanza, M. R. & Caneva, G. Natural biocides for the conservation of stone cultural heritage: A review. J Cult Herit 38, 271–286 (2019). 10.1016/j.culher.2019.01.005

12 Li, Q., Wu, C., He, J. & Zhang, B. Unraveling the microbiotas and key genetic contexts identified on stone heritage using illumina and nanopore sequencing platforms. Int Biodeter Biodegr 185, 105688 (2023). 10.1016/j.ibiod.2023.105688

13 Ding, X. et al. Microbiome characteristics and the key biochemical reactions identified on stone world cultural heritage under different climate conditions. Journal of Environmental Management 302, 114041 (2022). 10.1016/j.jenvman.2021.114041

14 Ennis, N. J., Dharumaduri, D., Bryce, J. G. & Tisa, L. S. Metagenome Across a Geochemical Gradient of Indian Stone Ruins Found at Historic Sites in Tamil Nadu, India. Microb Ecol 81, 385–395 (2021). 10.1007/s00248-020-01598-3

15 Louati, M. et al. Elucidating the ecological networks in stone-dwelling microbiomes. Environ Microbiol 22, 1467–1480 (2020). 10.1111/1462-2920.14700

16 Li, T., Cai, Y. & Ma, Q. Microbial Diversity on the Surface of Historical Monuments in Lingyan Temple, Jinan, China. Microb Ecol (2022). 10.1007/s00248-021-01955-w

17 Cattò, C. et al. Biofilm colonization of stone materials from an Australian outdoor sculpture: Importance of geometry and exposure. Journal of Environmental Management 339, 117948 (2023). 10.1016/j.jenvman.2023.117948

18 Zhou, J. & Ning, D. Stochastic Community Assembly: Does It Matter in Microbial Ecology? MMBR 81, e00002–00017 (2017). 10.1128/MMBR.00002-17

19 Dal Bello, M., Lee, H., Goyal, A. & Gore, J. Resource–diversity relationships in bacterial communities reflect the network structure of microbial metabolism. Nature Ecology & Evolution 5, 1424–1434 (2021). 10.1038/s41559-021-01535-8

20 Goldford, J. E. et al. Emergent simplicity in microbial community assembly. Science 361, 469–474 (2018). 10.1126/science.aat1168

21 Ning, D. et al. A quantitative framework reveals ecological drivers of grassland microbial community assembly in response to warming. Nat Commun 11, 4717 (2020). 10.1038/s41467-020-18560-z

22 Chase, J. M. Stochastic Community Assembly Causes Higher Biodiversity in More Productive Environments. Science 328, 1388–1391 (2010). 10.1126/science.1187820

23 Ofiţeru, I. D. et al. Combined niche and neutral effects in a microbial wastewater treatment community. PNAS 107, 15345–15350 (2010). 10.1073/pnas.1000604107

24 Chesson, P. Mechanisms of Maintenance of Species Diversity. Annual Review of Ecology, Evolution, and Systematics 31, 343–366 (2000). 10.1146/annurev.ecolsys.31.1.343

25 Fargione, J., Brown, C. S. & Tilman, D. Community assembly and invasion: An experimental test of neutral versus niche processes. PNAS 100, 8916–8920 (2003). 10.1073/pnas.1033107100

26 Chave, J. Neutral theory and community ecology. Ecol Lett 7, 241–253 (2004). 10.1111/j.1461-0248.2003.00566.x

27 Vega, N. M. & Gore, J. Stochastic assembly produces heterogeneous communities in the Caenorhabditis elegans intestine. PLOS Biology 15, e2000633 (2017). 10.1371/journal.pbio.2000633

28 Bashan, A. et al. Universality of human microbial dynamics. Nature 534, 259–262 (2016). 10.1038/nature18301

29 Ning, D. et al. Environmental stress mediates groundwater microbial community assembly. Nature Microbiology 9, 490–501 (2024). 10.1038/s41564-023-01573-x

30 He, Q. et al. Temperature and microbial interactions drive the deterministic assembly processes in sediments of hot springs. Sci Total Environ 772, 145465 (2021). 10.1016/j.scitotenv.2021.145465

31 Tolkkinen, M. et al. Multi-stressor impacts on fungal diversity and ecosystem functions in streams: natural vs. anthropogenic stress. Ecology 96, 672–683 (2015). 10.1890/14-0743.1

32 Huot, O. B. & Tamborindeguy, C. Drought stress affects olanum lycopersicum susceptibility to actericera cockerelli colonization. Entomologia Experimentalis et Applicata 165, 70–82 (2017). 10.1111/eea.12627

33 Luo, C., Lü, F., Shao, L. & He, P. Application of eco-compatible biochar in anaerobic digestion to relieve acid stress and promote the selective colonization of functional microbes. Water Res 68, 710–718 (2015). 10.1016/j.watres.2014.10.052

34 Bai, F. et al. Microbial biofilms on a giant monolithic statue of Buddha: The symbiosis of microorganisms and mosses and implications for bioweathering. Int Biodeter Biodegr 156 (2021). ARTN 105106 10.1016/j.ibiod.2020.105106

35 Zhu, C. et al. Analysis of the Microbiomes on Two Cultural Heritage Sites. Geomicrobiol J, 1–10 (2022). 10.1080/01490451.2022.2137604

36 Zhu, C. et al. Unveiling the dual role of biocolonization: a case study on the deterioration and preservation of sandstone monuments in Leshan Giant Buddha, China. World Journal of Microbiology and Biotechnology 41, 25 (2025). 10.1007/s11274-024-04237-y

37 Stegen, J. C., Lin, X., Konopka, A. E. & Fredrickson, J. K. Stochastic and deterministic assembly processes in subsurface microbial communities. ISME J 6, 1653–1664 (2012). 10.1038/ismej.2012.22

38 Stegen, J. C. et al. Quantifying community assembly processes and identifying features that impose them. ISME J 7, 2069–2079 (2013). 10.1038/ismej.2013.93

39 Levins, R. Evolution in Changing Environments Some Theoretical Explorations. (MPB-2). (Princeton University Press, 1968).

40 Sexton, J. P., Montiel, J., Shay, J. E., Stephens, M. R. & Slatyer, R. A. Evolution of Ecological Niche Breadth. Annual Review of Ecology, Evolution, and Systematics 48, 183–206 (2017). 10.1146/annurev-ecolsys-110316-023003

41 von Meijenfeldt, F. A. B., Hogeweg, P. & Dutilh, B. E. A social niche breadth score reveals niche range strategies of generalists and specialists. Nature Ecology & Evolution 7, 768–781 (2023). 10.1038/s41559-023-02027-7

42 Meng, H., Zhang, X., Katayama, Y., Ge, Q. & Gu, J.-D. Microbial diversity and composition of the Preah Vihear temple in Cambodia by high-throughput sequencing based on genomic DNA and RNA. Int Biodeter Biodegr 149 (2020).

43 Meng, S. et al. Community structures and biodeterioration processes of epilithic biofilms imply the significance of micro-environments. Sci Total Environ 876, 162665 (2023). 10.1016/j.scitotenv.2023.162665

44 Chen, X. et al. The Organisms on Rock Cultural Heritages: Growth and Weathering. Geoheritage 13 (2021). ARTN 5610.1007/s12371-021-00588-2

45 Escalas, A. et al. Microbial functional diversity: From concepts to applications. Ecology and Evolution 9, 12000–12016 (2019). 10.1002/ece3.5670

46 Prosser, J. I. et al. The role of ecological theory in microbial ecology. Nat Rev Microbiol 5, 384–392 (2007). 10.1038/nrmicro1643

47 Sterflinger, K. et al. Future directions and challenges in biodeterioration research on historic materials and cultural properties. Int Biodeter Biodegr 129, 10–12 (2018). 10.1016/j.ibiod.2017.12.007

48 Wu, Y. et al. Monitoring the Deterioration of Masonry Relics at a UNESCO World Heritage Site. KSCE Journal of Civil Engineering 25, 3097–3106 (2021). 10.1007/s12205-021-1716-z

49 Zhang, X. W., Ge, Q. Y., Zhu, Z. B., Deng, Y. M. & Gu, J. D. Microbiological community of the Royal Palace in Angkor Thom and Beng Mealea of Cambodia by Illumina sequencing based on 16S rRNA gene. Int Biodeter Biodegr 134, 127–135 (2018). 10.1016/j.ibiod.2018.06.018

50 Schröer, L., De Kock, T., Cnudde, V. & Boon, N. Differential colonization of microbial communities inhabiting Lede stone in the urban and rural environment. Sci Total Environ 733, 139339 (2020). 10.1016/j.scitotenv.2020.139339

51 Qian, Y., Liu, X., Hu, P., Gao, L. & Gu, J.-D. Identifying the major metabolic potentials of microbial-driven carbon, nitrogen and sulfur cycling on stone cultural heritage worldwide. Sci Total Environ 954, 176757 (2024). 10.1016/j.scitotenv.2024.176757

52 Grossi, C. E. M. et al. Methylobacterium sp. 2A Is a Plant Growth-Promoting Rhizobacteria That Has the Potential to Improve Potato Crop Yield Under Adverse Conditions. Frontiers in Plant Science 11 (2020). 10.3389/fpls.2020.00071

53 Ramond, P., Galand, P. E. & Logares, R. Microbial functional diversity and redundancy: moving forward. FEMS Microbiology Reviews 49, fuae031 (2025). 10.1093/femsre/fuae031

54 Tian, L. et al. Deciphering functional redundancy in the human microbiome. Nat Commun 11, 6217 (2020). 10.1038/s41467-020-19940-1

55 Li, H. et al. Food provisioning results in functional, but not compositional, convergence of the gut microbiomes of two wild Rhinopithecus species: Evidence of functional redundancy in the gut microbiome. Sci Total Environ 858, 159957 (2023). 10.1016/j.scitotenv.2022.159957

56 Moya, A. & Ferrer, M. Functional Redundancy-Induced Stability of Gut Microbiota Subjected to Disturbance. Trends Microbiol 24, 402–413 (2016). 10.1016/j.tim.2016.02.002

57 Tian, L. & Zhang, J. Dynamics of Rabies Epidemics in Vampire Bats. Complexity 2020, 7032451 (2020). 10.1155/2020/7032451

58 Talbot, J. M. et al. Endemism and functional convergence across the North American soil mycobiome. PNAS 111, 6341–6346 (2014). 10.1073/pnas.1402584111

59 Zeng, Q. et al. Stable functional structure despite high taxonomic variability across fungal communities in soils of old-growth montane forests. Microbiome 11, 217 (2023). 10.1186/s40168-023-01650-7

60 Chen, H. et al. Functional Redundancy in Soil Microbial Community Based on Metagenomics Across the Globe. Front Microbiol 13 (2022).

61 Zhou, J. et al. The diversity and ecological significance of microbial traits potentially involved in B12 biosynthesis in the global ocean. mLife 2, 416–427 (2023). 10.1002/mlf2.12095

62 Burke, C., Steinberg, P., Rusch, D., Kjelleberg, S. & Thomas, T. Bacterial community assembly based on functional genes rather than species. PNAS 108, 14288–14293 (2011). 10.1073/pnas.1101591108

63 Louca, S. et al. Function and functional redundancy in microbial systems. Nature Ecology & Evolution 2, 936–943 (2018). 10.1038/s41559-018-0519-1

64 Chikina, M., Robinson, J. D. & Clark, N. L. Hundreds of Genes Experienced Convergent Shifts in Selective Pressure in Marine Mammals. Molecular Biology and Evolution 33, 2182–2192 (2016). 10.1093/molbev/msw112

65 Winemiller, K. O., Fitzgerald, D. B., Bower, L. M. & Pianka, E. R. Functional traits, convergent evolution, and periodic tables of niches. Ecol Lett 18, 737–751 (2015). 10.1111/ele.12462

66 Nie, Y. et al. Species Divergence vs. Functional Convergence Characterizes Crude Oil Microbial Community Assembly. Front Microbiol 7 (2016).

67 Li, P. et al. Global diversity and biogeography of potential phytopathogenic fungi in a changing world. Nat Commun 14, 6482 (2023). 10.1038/s41467-023-42142-4

68 Gao, C. et al. Fungal community assembly in drought-stressed sorghum shows stochasticity, selection, and universal ecological dynamics. Nat Commun 11, 34 (2020). 10.1038/s41467-019-13913-9

69 Wang, B., Zhu, C., Hu, Y., Zhang, B. & Wang, J. Dynamics of microbial community composition during degradation of silks in burial environment. Sci Total Environ 883, 163694 (2023). 10.1016/j.scitotenv.2023.163694

70 Meng, H., Luo, L., Chan, H. W., Katayama, Y. & Gu, J.-D. Higher diversity and abundance of ammonia-oxidizing archaea than bacteria detected at the Bayon Temple of Angkor Thom in Cambodia. Int Biodeter Biodegr 115, 234–243 (2016). 10.1016/j.ibiod.2016.08.021

71 Puente-Sánchez, F., Pascual-García, A., Bastolla, U., Pedrós-Alió, C. & Tamames, J. Cross-biome microbial networks reveal functional redundancy and suggest genome reduction through functional complementarity. Communications Biology 7, 1046 (2024). 10.1038/s42003-024-06616-5

72 MacArthur, R. Species packing and competitive equilibrium for many species. Theoretical Population Biology 1, 1–11 (1970). 10.1016/0040-5809(70)90039-0

73 Advani, M., Bunin, G. & Mehta, P. Statistical physics of community ecology: a cavity solution to MacArthur’s consumer resource model. Journal of Statistical Mechanics: Theory and Experiment 2018, 033406 (2018). 10.1088/1742-5468/aab04e

74 Marsland, R., III et al. Available energy fluxes drive a transition in the diversity, stability, and functional structure of microbial communities. PLOS Computational Biology 15, e1006793 (2019). 10.1371/journal.pcbi.1006793

75 Sriswasdi, S., Yang, C.-c. & Iwasaki, W. Generalist species drive microbial dispersion and evolution. Nat Commun 8, 1162 (2017). 10.1038/s41467-017-01265-1

76 Wang, B., Qi, M., Ma, Y., Zhang, B. & Hu, Y. Microbiome Diversity and Cellulose Decomposition Processes by Microorganisms on the Ancient Wooden Seawall of Qiantang River of Hangzhou, China. Microb Ecol 86, 2109–2119 (2023). 10.1007/s00248-023-02221-x

77 Wang, B., Zhu, C., Wang, B., Zhang, B. & Hu, Y. Analysis of the biocorrosion community from ancient wooden constructions at Tianluoshan (7000–6300 cal BP), Zhejiang Province, China. Heritage Science 12, 189 (2024). 10.1186/s40494-024-01304-3

78 Zhu, C. et al. Application and evaluation of a new blend of biocides for biological control on cultural heritages. Int Biodeter Biodegr 178, 105569 (2023). 10.1016/j.ibiod.2023.105569

79 Li, J. et al. Changes in soil microbial communities at Jinsha earthen site are associated with earthen site deterioration. BMC Microbiology 20, 147 (2020). 10.1186/s12866-020-01836-1

80 Wang, X., Wang, B., Hu, Y., Zhang, Z. & Zhang, B. Activity-based protein profiling technology reveals malate dehydrogenase as the target protein of cinnamaldehyde against Aspergillus niger. Int J Food Microbiol 417, 110685 (2024). 10.1016/j.ijfoodmicro.2024.110685

81 Logue, J. B. et al. Experimental insights into the importance of aquatic bacterial community composition to the degradation of dissolved organic matter. ISME J 10, 533–545 (2016). 10.1038/ismej.2015.131

82 Chen, T. et al. The Genome Sequence Archive Family: Toward Explosive Data Growth and Diverse Data Types. Genomics, Proteomics & Bioinformatics 19, 578–583 (2021). 10.1016/j.gpb.2021.08.001

83 Members, C.-N. & Partners. Database Resources of the National Genomics Data Center, China National Center for Bioinformation in 2022. Nucleic Acids Research 50, D27–D38 (2022). 10.1093/nar/gkab951

84 Callahan, B. J. et al. DADA2: High-resolution sample inference from Illumina amplicon data. Nat Methods 13, 581–583 (2016). 10.1038/nmeth.3869

85 Bolyen, E. et al. Reproducible, interactive, scalable and extensible microbiome data science using QIIME 2. Nat Biotechnol 37, 852–857 (2019). 10.1038/s41587-019-0209-9

86 Wang, Q., Garrity, G. M., Tiedje, J. M. & Cole, J. R. Nave Bayesian Classifier for Rapid Assignment of rRNA Sequences into the New Bacterial Taxonomy. Appl Environ Microbiol 73, 5261 (2007).

87 Pruesse, E. et al. SILVA: a comprehensive online resource for quality checked and aligned ribosomal RNA sequence data compatible with ARB. Nucleic Acids Res 35, 7188–7196 (2007). 10.1093/nar/gkm864

88 Nilsson, R. H. et al. The UNITE database for molecular identification of fungi: handling dark taxa and parallel taxonomic classifications. Nucleic Acids Research 47, D259–D264 (2019). 10.1093/nar/gky1022

89 Fu, L., Niu, B., Zhu, Z., Wu, S. & Li, W. CD-HIT: accelerated for clustering the next-generation sequencing data. Bioinformatics 28, 3150–3152 (2012). 10.1093/bioinformatics/bts565

90 Langmead, B. & Salzberg, S. L. Fast gapped-read alignment with Bowtie 2. Nat Methods 9, 357–359 (2012). 10.1038/nmeth.1923

91 Buchfink, B., Xie, C. & Huson, D. H. Fast and sensitive protein alignment using DIAMOND. Nat Methods 12, 59–60 (2015). 10.1038/nmeth.3176

92 Tu, Q., Lin, L., Cheng, L., Deng, Y. & He, Z. NCycDB: a curated integrative database for fast and accurate metagenomic profiling of nitrogen cycling genes. Bioinformatics 35, 1040–1048 (2019). 10.1093/bioinformatics/bty741

93 Wickham, H. ggplot2. WIREs Computational Statistics 3, 180–185 (2011). 10.1002/wics.147

94 Dixon, P. VEGAN, a package of R functions for community ecology. Journal of Vegetation Science 14, 927–930 (2003). 10.1111/j.1654-1103.2003.tb02228.x

95 Kembel, S. W. et al. Picante: R tools for integrating phylogenies and ecology. Bioinformatics 26, 1463–1464 (2010). 10.1093/bioinformatics/btq166

96 Nguyen, L.-T., Schmidt, H. A., von Haeseler, A. & Minh, B. Q. IQ-TREE: A Fast and Effective Stochastic Algorithm for Estimating Maximum-Likelihood Phylogenies. Molecular Biology and Evolution 32, 268–274 (2015). 10.1093/molbev/msu300

97 Zhang, J. M., KP. spaa: An R package for computing species association and niche overlap. . Research Progress of Biodiversity Conservation in China (in Chinese) **X**, 165–174 (2014).

98 Pandit, S. N., Kolasa, J. & Cottenie, K. Contrasts between habitat generalists and specialists: an empirical extension to the basic metacommunity framework. Ecology 90, 2253–2262 (2009). 10.1890/08-0851.1

99 Hao, X. et al. Effects on community composition and function Pinus massoniana infected by Bursaphelenchus xylophilus. BMC Microbiology 22, 157 (2022). 10.1186/s12866-022-02569-z

100 Wen, T. et al. ggClusterNet: An R package for microbiome network analysis and modularity-based multiple network layouts. iMeta 1, e32 (2022). 10.1002/imt2.32

101 ggraph: An Implementation of Grammar of Graphics for Graphs and Networks (2025).

102 Csardi, G. & Nepusz, T. The igraph software package for complex network research. InterJournal, Complex Systems 1695 (2006).

